# Structural and functional specializations of human fast spiking neurons support fast cortical signaling

**DOI:** 10.1101/2022.11.29.518193

**Authors:** René Wilbers, Anna A. Galakhova, Stan L.W. Driessens, Tim S. Heistek, Verjinia D. Metodieva, Jim Hagemann, Djai B. Heyer, Eline J. Mertens, Suixin Deng, Sander Idema, Philip C. de Witt Hamer, David P. Noske, Paul van Schie, Ivar Kommers, Guoming Luan, Tianfu Li, Yousheng Shu, Christiaan P.J. de Kock, Huibert D. Mansvelder, Natalia A. Goriounova

## Abstract

Fast spiking interneurons (FSINs) provide fast inhibition that synchronizes neuronal activity and is critical for cognitive function. Fast synchronization frequencies are evolutionary conserved in the expanded human neocortex, despite larger neuron-to-neuron distances that challenge fast input-output transfer functions of FSINs. Here, we test in human neurons from neurosurgery tissue which mechanistic specializations of human FSINs explain their fast-signaling properties in human cortex. With morphological reconstructions, multi-patch recordings, and biophysical modeling we find that despite three-fold longer dendritic path, human FSINs maintain fast inhibition between connected pyramidal neurons through several mechanisms: stronger synapse strength of excitatory inputs, larger dendrite diameter with reduced complexity, faster AP initiation, and faster and larger inhibitory output, while Na^+^ current activation/inactivation properties are similar. These adaptations underlie short input-output delays in fast inhibition of human pyramidal neurons through FSINs, explaining how cortical synchronization frequencies are conserved despite expanded and sparse network topology of human cortex.

**Teaser/one-sentence summary:** Specializations of fast spiking human neurons ensure fast signaling in human cortex.

## Introduction

Information flow in the mammalian neocortex is shaped through canonical motifs of connected excitatory pyramidal projection neurons and a diverse population of inhibitory GABAergic interneurons (*1, 2*). Incoming activity to pyramidal neurons is regulated through feedforward, feedback, lateral and disinhibitory motifs before it is passed on to downstream target brain areas. In the rodent cortex, distinct but well-defined, types of interneurons make up these inhibitory motifs: fast spiking parvalbumin+ cells provide fast feedforward, feedback and fast lateral inhibition of pyramidal neuron somatodendritic regions, somatostatin+ interneurons generate delayed lateral inhibition on dendrites, while VIP+ interneurons disinhibit pyramidal neurons by inhibiting other types of interneurons (*1, 2*). Whether these fundamental cortical processing motifs operate in the strongly expanded human neocortex is poorly understood.

Fast spiking interneurons (FSINs) provide fast inhibitory motifs in rodent neocortex that synchronize neuronal activity at gamma frequencies underlying cognitive and sensory function (*2, 3*). These neurons provide fast, reliable, strong and precise inhibition of target cells (*2, 3*), and have been found in the cortices of mice, rats, marmosets, monkeys and humans, with characteristic fast spiking phenotypes (*4*–*7*). FSINs project primarily to pyramidal neurons and other FSINs (*8, 9*) and possess several specializations to support fast input-output function. Fast dendritic AMPA receptors in combination with fast potassium channels (Kv3) result in short excitatory postsynaptic potentials (EPSPs) and stable coupling of incoming input (EPSPs) to neuronal output - action potential (AP) firing (*10*). A high resting membrane potential in the soma combined with a high density of Na^+^ channels in the axon (*3*) ensures that FS neuron can quickly reach threshold to initiate an AP. Moreover, axons of FSINs are partly myelinated to support fast conduction (*11, 12*) and synaptic boutons are equipped with ultra-fast release machinery (*13*). These properties result in extremely fast disynaptic inhibition where one pyramidal neuron inhibits a neighboring pyramidal neuron via an intermediate FSIN with short delays of 3-6 ms in both human and rodent cortex (*9, 14, 15*). In contrast, disynaptic inhibition loops through somatostatin-positive Martinotti neurons have a delay of approximately 100 ms (*16, 17*). Fast loops help to narrow the temporal window for neuronal integration and improve temporal resolution of neuronal signaling, but they require very fast conversion of inputs into outputs in FSINs.

In humans, the evolutionary expansion of cortex and especially its upper cortical layers (layers 2 and 3, L2/L3), is accompanied by a three-fold increase in the size and complexity of human pyramidal neurons and their dendrites, while the neuronal density is lower (*18*–*20*). Human cortical cytoarchitecture with sparse but larger neurons could have dramatic consequences for the fast operation of FSINs, since increased neurite length could potentially slow down their input-output processing in two ways. Firstly, if human FSIN dendrites are longer, the excitatory inputs need to travel a longer dendritic path. This can result in reduced EPSP amplitude and kinetics through dendritic filtering (*21*), potentially delaying conversion of synaptic input to AP. Secondly, if human FSIN axonal paths are also longer because of increased distances between neurons in the human cortex, it may cause an extensive delay in synaptic output due to the delay caused by conduction along the axon. Therefore, if dendritic and axonal paths of human FSINs are similarly elongated as observed in human pyramidal neurons (*18*), this might result in derogation of their fast function. Despite the expanded and sparse network topology of human neocortex (*18*–*20*), the gamma frequency range of synchronized brain activity is preserved across mammalian species (*22*). This may suggest that adaptations in human cortex exist to accommodate larger neuronal structures. It is unclear which biophysical mechanisms enable human FS neurons to achieve fast input-output function.

Here, we addressed this question in nonpathological human cortical tissue that was resected as part of neurosurgical treatment to gain access to deeper lying disease focus (epilepsy or tumor). With a combination of detailed morphological reconstructions of FSINs in human and mouse, multi-patch electrophysiological recordings of connected FSINs and pyramidal neurons and computational models we identify several biophysical specializations in human FSINS that preserve a fast input-output transfer function.

## Results

### Human FSIN dendrites have elongated path lengths with reduced complexity

In human cortex, pyramidal neurons have more elaborate dendritic trees and are distributed less densely throughout the layers, resulting in large neuron-to-neuron distances compared with other primates or rodents (*18*), but human interneuron morphology is less well documented. We first asked how dendritic structure of human L2/L3 FSINs compares with mouse FSIN dendrites, and made dendritic reconstructions of 16 human and 43 mouse FSINs from layers 2 and 3 of various cortical areas (human: 13 temporal cortex, 3 frontal cortex; mouse: 10 temporal association cortex, 33 primary visual cortex). We quantified several morphological parameters that describe the dendritic structure of the neuron (Fig 1A, Fig S1): the sum length of all dendrites per neuron (total dendritic length, Fig 1B), number of dendritic branch points (Fig 1C), number of dendrites growing directly from soma (Fig 1D), the thickness of each dendrite segment (Fig 1E), maximal path length from soma to the tip of the longest dendrite (Fig 1F), the non-terminal dendritic segment lengths (length of dendrite between two consecutive branching points, excluding terminal segments, Fig 1G) and terminal segments (Fig 1H). We find that human FSIN dendrites were generally longer: the median (Q1-Q3) total dendritic length was 3.5 (3.1-4.2) mm in humans and 2.1 (1.7-2.6) mm in mice (Fig. 1B). Interestingly, the increase in dendrite length was accompanied by a decrease in dendritic complexity, as human FSINs had fewer branches (Fig. 1C, mean±SD number of branch points, human: 12.8±6.1, mouse: 17.7±6.4) and fewer dendrites stemming from the soma (Fig. 1D, median [Q1-Q3], human: 5 [4-6] mouse: 7 [6-7]). As dendritic filtering would primarily be influenced by dendritic diameter and the distance excitatory postsynaptic potentials (EPSPs) must travel along a dendrite, we analyzed diameters and maximal path lengths of dendrites. The human dendrites were thicker over the entire range of branch orders (Fig. 1E) and their path length from dendritic tip to soma in human FSINs was about two times longer (Fig. 1F, mean±SD, human: 248±62, mouse: 123±23 μm). Next, we asked which dendritic segments were contributing most to longer human dendrites. We find that exclusively terminal segments are responsible for the increase in path length (Fig. 1G-H; mean±SD non-terminal segment length, human: 17.2±11.1, mouse: 25.7±9.4 μm; median [Q1-Q3] terminal segment length, human: 216 [152-241], mouse: 72 [62-83] μm). Although some differences were observed between mouse FSIN morphologies from temporal and visual areas, when we compared human data with mouse dataset from temporal and visual areas separately, the conclusions remained unchanged (Fig. S1). Thus, dendrites of human FSINs have a different structure compared to mouse, they are thicker, less numerous and have longer path lengths that can solely be attributed to long terminal segments.

**Figure 1.**
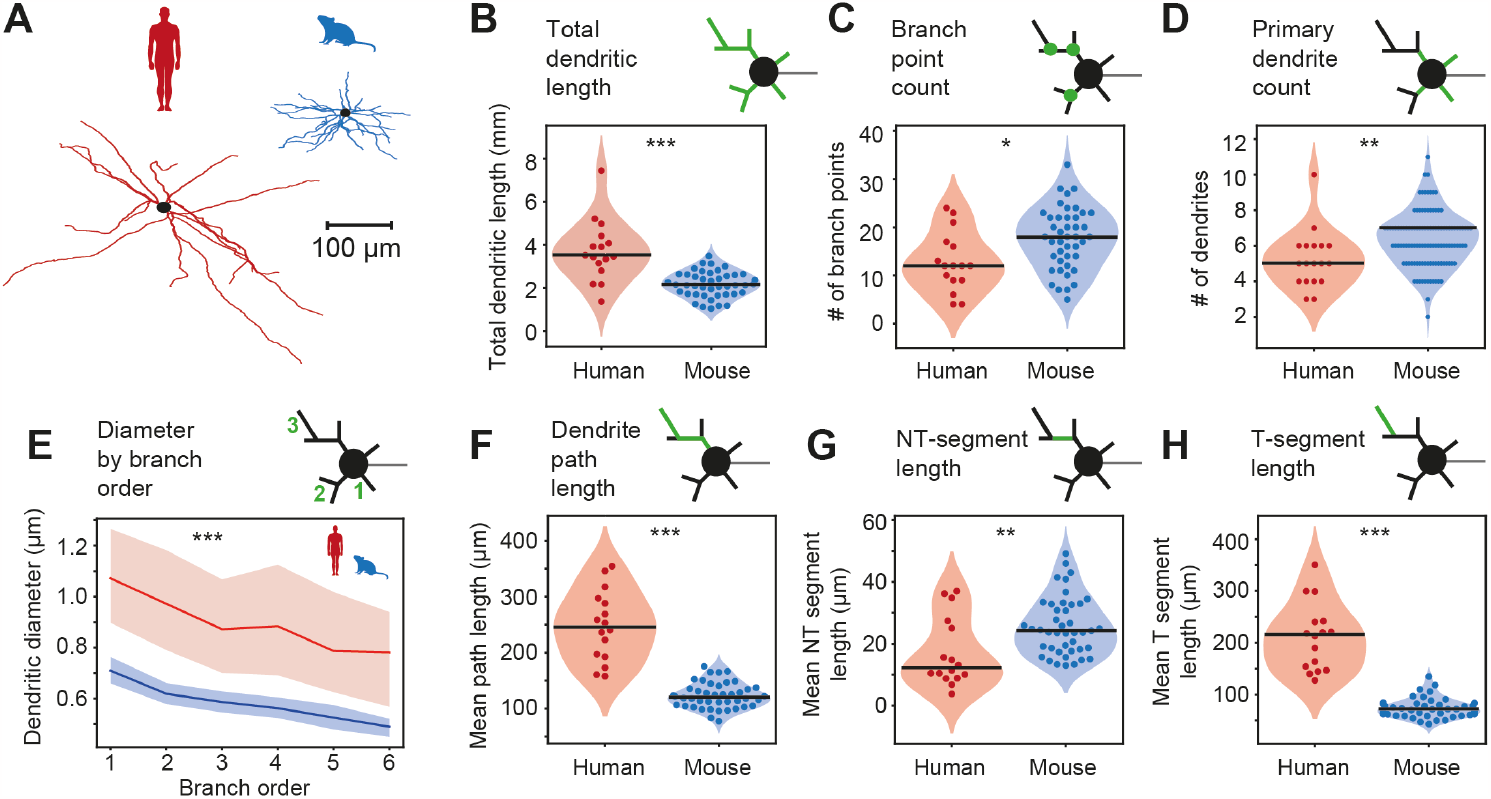
Human FSIN dendrites have elongated path lengths with reduced complexity. (A) Examples of dendritic reconstructions of L2/L3 FSINs from human and mouse cortices. (B) Total dendritic length. ***p<10^-4^, Wilcoxon rank sum (WRS) test. (C) Number of branch points *p=0.01, t-test. (D) Number of primary dendrites originating from the soma. **p=0.004, WRS test. (E) Dendritic diameters over branch orders. ***p<10^-12^, linear regression model (species effect). (F) Mean dendritic path length from terminal endpoints to soma. ***p<10^-6^, t-test. (G) Mean length of non-terminal (NT) segments. **p=0.0045, t-test. (H) Mean length of terminal segments. ***p<10^-8^, WRS test. Human: n=16 FSINs; mouse n=43 FSINs.

### Excitatory inputs have similar strength and kinetics in human and mouse FSINs

As dendritic morphology shapes dendritic filtering of incoming EPSPs, we wondered whether the amplitude and speed of incoming EPSPs was different in human FSINs. To test this, we investigated simultaneous recordings of synaptically connected pyramidal neurons to FSINs in L2/L3 of human and mouse cortices (Fig. 2A). Surprisingly, despite the elongated dendrites in human FSINs, incoming unitary EPSPs from pyramidal neurons were comparable in strength across FSINs from species: there were no significant differences in unitary EPSP amplitude (Fig. 2B, mean ± SD: human 1.65 ± 1.59 mV; mouse 1.22 ± 1.03 mV). Furthermore, as dendritic filtering does not only affect EPSP amplitude, but also the shape of incoming inputs, we analyzed the onset and decay kinetics of the averaged EPSP traces. We found no differences between the species: EPSPs had similar onset latency (Fig. 2C, mean ± SD: human 1.31 ± 0.50 ms; mouse 1.14 ± 0.36 ms), rise time (Fig. 2D, mean ± SD: human 2.05 ± 1.36 ms; mouse 1.84 ± 0.65 ms) and time constant of decay (Fig. 2E, mean ± SD: human 17.6 ± 13.4 ms; mouse 17.2 ± 18.5 ms). These results indicate that the morpho-physiological characteristics of human FSINs result in a similar amount of dendritic filtering of EPSPs even though their dendrites are over two times longer.

**Figure 2.**
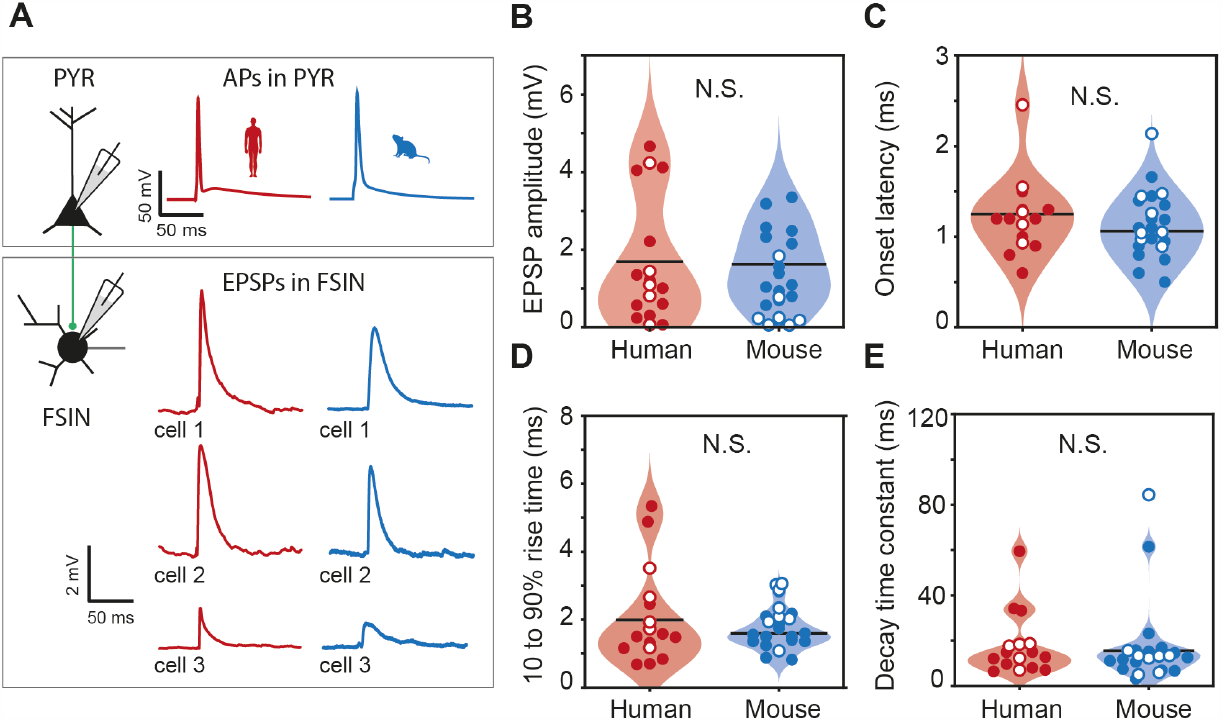
Unitary EPSP strength and kinetics are similar in human and mouse FSINs. **(A)** Schematic representation and example traces from 3 human and 3 mouse recordings of pyramidal-FSIN pairs, where single APs in a pyramidal neuron evoked unitary EPSPs in a FSIN. (B-F) EPSP parameters extracted from average trace of 5-10 traces (data points), their distributions (violins) and median values (black horizontal lines) are shown for the data recorded in this study (filled circles) and extracted from AIBS database (open circles). Sample size: human n=17 connected pairs: 12 our lab (filled circles), 5 from AIBS (open circles); mouse n=23 connected pairs: 15 our lab (filled circles), 8 from AIBS (open circles). (B) EPSP amplitude p=0.46, Mann-Whitney-U (MWU) test. (C) EPSP onset latency p=0.23, MWU test. (D) 10 to 90% rise time p=0.54, MWU test. (F) Time constant of decay p=0.48, MWU test. N.S.: nonsignificant.

### Conserved Na^+^ current kinetics in human and mouse FSINs

To understand how different dendritic length results in similar EPSP shape and amplitude across species we created detailed conductance-based computational FSIN models with realistic morphology. The model should be able to predict EPSP characteristics as well as AP timing, which ultimately determines the delay of synaptic output. However, as Na^+^ channels play a crucial role in AP initiation we first asked whether the Na^+^ current activation and inactivation properties are different between human and mouse FSINs. To this end, we recorded Na^+^ currents in nucleated patches of mouse and human L2/L3 FSINs and characterized the voltage dependent properties of these currents. First, we obtained inactivation and activation curves by determining maximal conductance during different pre-pulse and pulse voltages respectively (Fig. 3A). Fitting Boltzmann curves to the data showed that the half-voltages for inactivation (Fig. 3B, median [Q1-Q3], human: -25.1 [-27.8 - -16.7]; mouse -25.6 [-29.0 - -21.8] mV) and activation, (Fig. 3B, median [Q1-Q3], human: -48.9 [-56.2 - -36.4]; mouse -51.6 [-53.9 - -45.9] mV) were not significantly different between species. As the speed of AP initiation is also highly dependent on how rapidly Na^+^ channels activate, we next measured the time constant of activation by fitting an exponential to the activation phase of the current. The time constant of activation was highly dependent on voltage but was not different between mouse and human FSINs (Fig. 3C). As AP initiation is also influenced by the amount of functionally available channels we further determined the inactivation time constants. The time constant of inactivation was similar for both species across voltages (Fig. 3D). Next, we fully inactivated Na^+^ channels and used pulses after various recovery voltages and delays to assess the time course and voltage-dependence of Na^+^ current recovery. We fitted exponentials to the amplitudes of evoked Na^+^ currents to determine their time constant and found that both mouse and human FSINs recovered from inactivation with similar time constants (Fig. 3E). These results indicate that it is unlikely that any differences in AP timing between mouse and human FSINs are caused by differences in Na^+^ current kinetics, and other mechanisms might be involved.

**Figure 3.**
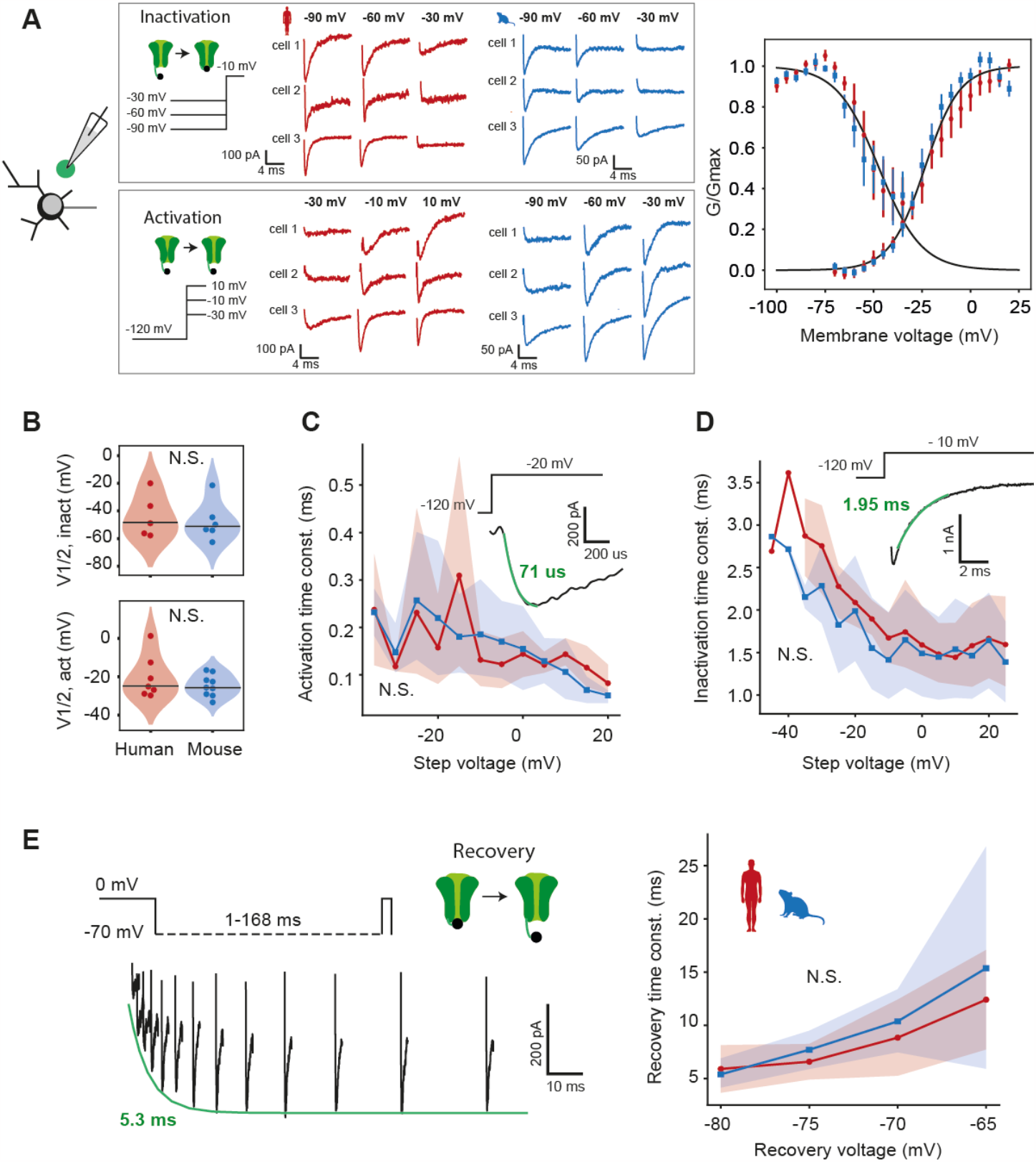
Conserved somatic Na^+^ current properties in human and mouse L2/L3 FSINs. (A) Somatic Na^+^ current activation and inactivation properties are similar in mouse and human FSINs. Somatic Na^+^ currents were recorded in nucleated patch recordings, example Na^+^ currents for 3 mouse and 3 human neurons at different pre-pulse voltages in inactivation protocol and at different activation voltages in activation protocol are shown in gray frames. Inactivation and activation curves (Mean±SEM) from human (red) and mouse (blue) FSINs are shown right Black lines: Boltzmann fits. (B) Half-voltages of inactivation (top) and activation (bottom) from Boltzmann fits to individual FSINs. (C) Time constant of activation across voltages. Inset: example trace from a human nucleated patch recording. N.S.: p=0.90, linear regression model (species effect). (D) Time constant of inactivation across voltages. Inset: example trace from human patch. N.S.: p=0.10, linear regression model (species effect). (E) Left: example trace in a human FSIN showing the recovery protocol. Right: Time constant of recovery at -80 to -65 mV. N.S.: p=0.62, linear regression model (species effect). Sample size: human n=7 recordings, mouse n=9.

### Mechanisms of fast AP responses to distal synaptic inputs

The longer dendrites of human FSINs could potentially lead to longer delays of synaptic inputs from dendrites to soma, but we observed that unitary EPSPs from presynaptic pyramidal neurons did not have larger onset-latencies in human FSINs. To understand how individual structural and physiological properties of human FSINs may contribute to fast input (EPSP) to output (AP) conversion, we built a detailed conductance-based Hodgkin-Huxley computational model. To keep the feature space constant, the model neurons had artificial morphologies based on experimentally observed morphological features (Fig. 1). This resulted in a mouse-like and a human-like artificial morphologies (Fig. 4A). After fitting the densities of Na^+^, K^+^, HCN and leak conductances for each cellular compartment we obtained a stable model with characteristic FS responses to long current steps (Fig. 4B). We kept these conductances the same for mouse and human model neurons, that only differed in their dendritic structure (model parameters are shown in Tables 1 and 2).

**Table 1.** Feature objectives for model optimization.

| Protocol | Feature | Objective | SD |
| --- | --- | --- | --- |
| **100 ms, 200 pA step** | Time to first spike | 10 ms | 1 ms |
|  | AP1 peak | 30 mV | 5 mV |
|  | AP1 halfwidth | 0.3 ms | 0.05 ms |
|  | AP1 rise speed | 350 mV/ms | 70 mV/ms |
|  | AP1 fall speed | -180 mV/ms | 40 mV/ms |
|  | Absolute AHP | -60 mV | 10 mV |
|  | Spike count | 10 | 5 |
|  | Peak AP1 – Peak AP2 | 0 mV | 4 mV |
| **100 ms, -100 pA Step** | Voltage deflection (steady state) | 10 mV | 3 mV |
|  | Decay time constant after stim | 12 ms | 4 ms |
|  | Sag ratio | 0.1 | 0.04 |
| **Short pulse: 800 pA, 4 ms** | Spike count (soma) | 1 | 0.0001 |
|  | Spike count (AIS) | 1 | 0.0001 |
|  | Spike count (distal axon) | 1 | 0.0001 |
|  | Time to first spike (soma) | 2.9 ms | 0.1 ms |
|  | Time to first spike (AIS) – time to first spike (soma) | 0.05 ms | 0.005 ms |
|  | Time to first spike (distal axon) – time to first spike (soma) | 0.45 ms | 0.15 ms |
| **Synapse activation (4 x 0.01 nS at 108 um from Soma)** | Spike count (soma) | 1 | 0.0001 |
|  | Spike count (AIS) | 1 | 0.0001 |
|  | Time to first spike (AIS) – time to first spike (soma) | 0.05 ms | 0.005 ms |

**Table 2.** Optimized parameter values used in the final model.

| Parameter | Location | Value |
| --- | --- | --- |
| Passive conductance density | Dendrite | 0.003 pS*$\mu\text{m}^{-2}$ |
| | Soma | 30.545 pS*$\mu\text{m}^{-2}$ |
| | Axon | 9.663 pS*$\mu\text{m}^{-2}$ |
| Na<sup>+</sup> conductance density | Dendrite | 89.308 pS*$\mu\text{m}^{-2}$ |
| | Soma | 405.812 pS*$\mu\text{m}^{-2}$ |
| | Axon | 1728.496 pS*$\mu\text{m}^{-2}$ |
| K<sup>+</sup> conductance density | Dendrite | 54.240 pS*$\mu\text{m}^{-2}$ |
| | Soma | 505.077 pS*$\mu\text{m}^{-2}$ |
| | Axon | 161.576 pS*$\mu\text{m}^{-2}$ |
| HCN conductance density | Dendrite | 8.367 pS*$\mu\text{m}^{-2}$ |
| | Soma | 3.716 pS*$\mu\text{m}^{-2}$ |
| | Axon | 6.722 pS*$\mu\text{m}^{-2}$ |
| Left shift of Na<sup>+</sup> channels | Axon | -0.62 mV |

**Figure 4.**
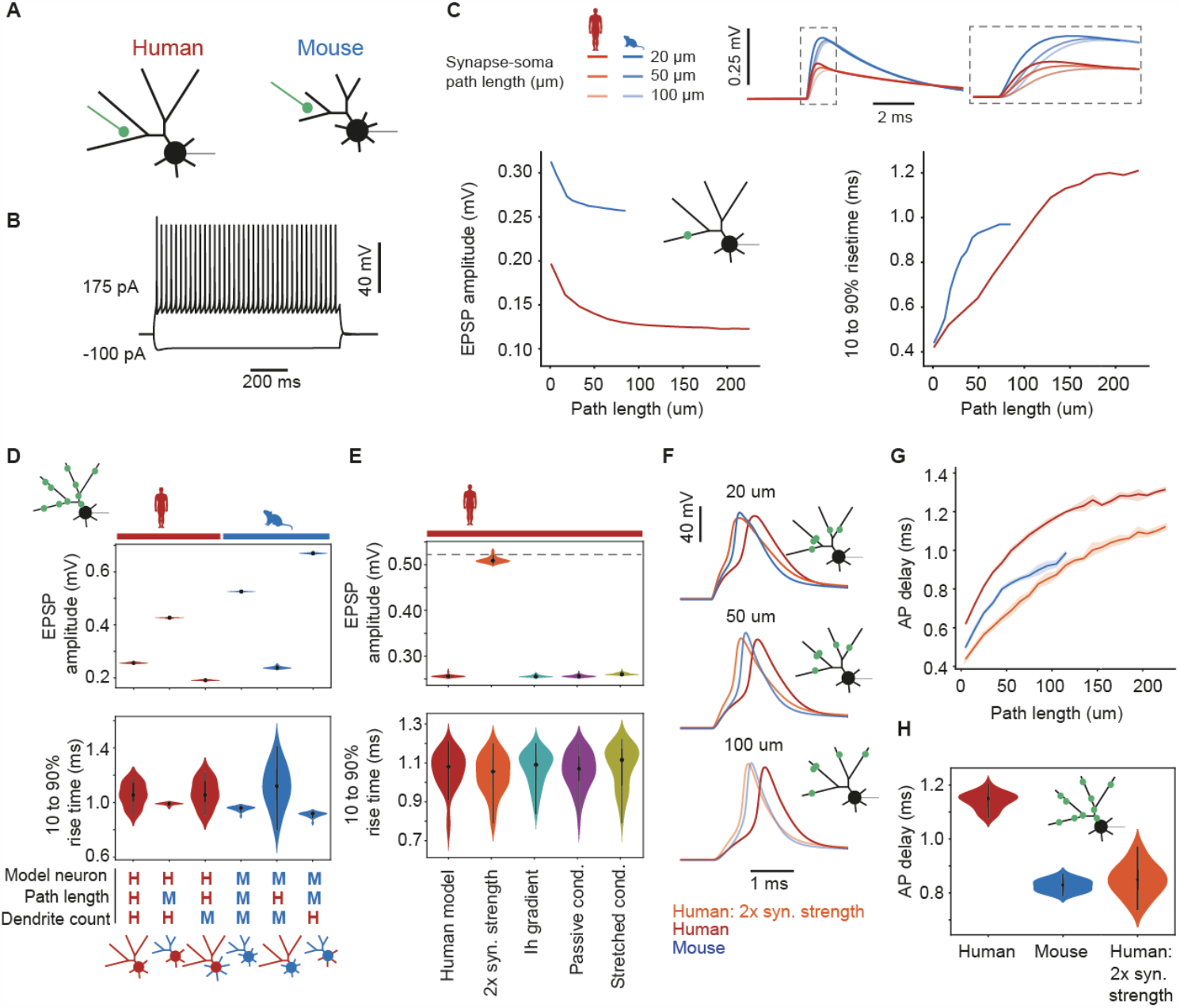
Human dendritic morphology combined with increased synapse size is sufficient to allow fast responses to distal excitatory inputs to FSINs. (A) Schematic representation of human and mouse FSIN models based on dendrite count, dendrite diameter, terminal segment length, and non-terminal segment length. (B) Typical FSIN responses of the model to simulated current injection. (C) Somatic EPSP amplitude decrease and EPSP rise time increase as a function of synapse distance in models based on human and mouse morphologies. Example EPSP traces of human and mouse FSIN models to synaptic stimulations at different distances from soma are shown above. (D) Amplitudes and rise times of somatic EPSPs generated in human, mouse and hybrid models based on morphological parameters; from left to right: full human model, human model with mouse number of primary dendrites originating from soma (7 dendrites), human model with the dendritic path length from mouse neurons, full mouse model, mouse model with dendritic path length from human FSINs, mouse model with human number of primary dendrites originating from soma (5 dendrites). (E) Amplitudes and rise times of somatic EPSPs generated in human models with an addition of one of the following electrophysiological parameters, from left to right: human model, human model with a two-fold increase of synapse strength, with added Ih current gradient, with added passive conductance gradient, with added stretched conductance. (F) AP traces of APs generated by synaptic inputs at different synaptic distances in the mouse, human and human + two-fold increase in synapse strength models. (G) Summary data of AP delays in the three models generated by inputs at different synaptic distances (H) Stimulation of synaptic inputs at random locations leads to longer AP delays in the human model, that is rescued by a two-fold increase in synaptic strength.

Next, we asked how EPSPs arriving at the soma are affected by dendritic morphology and the distance of the dendritic synaptic input site from the soma. When we activated synapses at different dendritic locations we noticed that both models showed a distance-dependent decrease in amplitude and increase in risetime, as expected from dendritic filtering (Fig. 4C). However, human model neurons showed smaller EPSP amplitudes and shorter rise times when measured at the soma in response to inputs at the same dendritic distance as in the mouse model (Fig. 4C). As real EPSPs are not generated by a single synapse we activated several synapses simultaneously at random dendritic locations: at 20 different random configurations where each configuration had 10 randomly distributed synapse locations. Similar to the activation of single synapse, in model neurons based on human dendritic morphology, the combined EPSPs had consistently smaller amplitudes and the variability of the rise times was considerably increased (Fig. 4D). Next, to understand which morphological features cause this effect we generated several hybrid morphologies, where we changed only one of the dendritic parameters between human and mouse models: path length or the number of dendrites. First, when we elongated the path and segment lengths in the mouse model to match lengths observed in human FSINs, the EPSP amplitude dropped (Fig. 4D). Conversely, shortening path lengths in the human model resulted in a higher EPSP amplitude. However, the effect of path length on EPSP amplitude was much smaller in the human model than in the mouse model, demonstrating that other morphological features counteract the reduction in amplitude that comes with long dendrites. When we increased the number of dendrites in the human model to 7 as was observed in mouse FSINs, this manipulation strongly reduced EPSP amplitude compared with the full human model (Fig 4D). This suggests that the lower number of dendrites in human FSINs compensates for longer dendrites and reduces dendritic filtering (Fig 4D). In addition, given the strong relationship between dendritic path length and EPSP rise time (Fig. 4C), the increase in rise time in the human model was only moderate (Fig. 4D). Furthermore, the rise time of the human model was slightly faster and less variable than the rise time in a mouse model with long dendritic path lengths (Fig. 4D). Thus, our results from models based on realistic morphological parameters show that longer human FSIN dendrites might lead to increased dendritic filtering and smaller EPSP amplitudes, while a lower number of dendrites counteracts this effect.

Our findings from mouse and human models were not consistent with our experimental results, where we found that EPSPs were similar in size and kinetics between species. In contrast, EPSP amplitude was two times lower in models based on human morphology. Hence, we hypothesized that there must be other physiological factors at play to further counteract EPSP filtering. Several physiological mechanisms were previously suggested to accelerate and boost EPSPs in human dendrites: 4-fold increased size of excitatory synapse on human FSINs (measured as the number of docked vesicles and functional release sites) (*7*), gradient of HCN channels (*21, 23*), a gradient in leak conductance (*24*) and stretched conductance (*25, 26*). We set out to test which of these mechanisms has the strongest effect on reducing dendritic filtering in the human model. We find that the only mechanism that could rescue the somatic EPSP amplitude to match the EPSP amplitude of the mouse model was the two-fold increase in excitatory synapse size (Fig. 4E). This indicates that, in order to preserve somatic EPSP amplitude as we have experimentally observed (Fig. 2B), a larger synaptic size is critical in human FSINs.

To produce output, FSINs need to integrate the synaptic inputs and generate AP. Fast input to output conversion requires fast AP generation and timing. We tested how FSIN dendritic morphology affected AP timing in response to synaptic input in human and mouse models that either only differed in morphology or where the human FSIN model had increased synapse size. We activated 20 synapses at different path lengths in these three models and measured delays to AP peak. We find that in all three models (mouse, human, and human + two-fold increased synaptic strength) more distal inputs led to longer AP delays (Fig. 4F, G). The AP delays were consistently longer in the model based on human morphology. However, in human model with larger synaptic strength the AP timing was the fastest (Fig. 4F-H). Our findings demonstrate that the stronger excitatory synapses and fewer dendrites in human FSINs are sufficient to boost EPSP transfer to soma and compensate for the longer human dendrites. These morphological and physiological specializations speed up generation of APs in response to excitatory synaptic inputs and improve input-output function of human FSINs.

### Human FS APs show fast onset kinetics facilitated by long dendritic length

The speed of FSIN output is not only dependent on the size and shape of EPSPs, but also on how fast APs can be generated by the neurons. To study AP generation, we experimentally recorded APs from human and mouse FSINs. To quantify AP onset kinetics, we analyzed AP waveform, derivative and phase plane in FSINs from both species (Fig. 5A, 10 human and 12 mouse FSINs). AP threshold, defined as the voltage at which the slope reached 5% of the maximum slope, was significantly lower in human FSINs (mean±SD, human -42.2 ± 3.1 mV, mouse -36.0 ± 3.5 mV, Fig. 5B). Furthermore, the initial slope in the phase plane, or ‘onset rapidity’, was significantly steeper (human 41.9 ± 4.8 ms^-1^, mouse 31.8 ± 6.3 ms^-1^), meaning that the AP initiates in human FSINs more suddenly rather than gradually. Furthermore, the overall AP speed in human FSINs was faster: human APs had faster rising speed (human 412 ± 115 mV/ms, mouse 268 ± 77 mV/ms) as well as falling speed (human 191 ± 65 mV/ms, mouse 123 ± 49 mV/ms). These experimental results support our findings from modeling and indicate that human FSIN neurons are able to respond faster and generate APs with fast kinetics and onsets.

**Figure 5.**
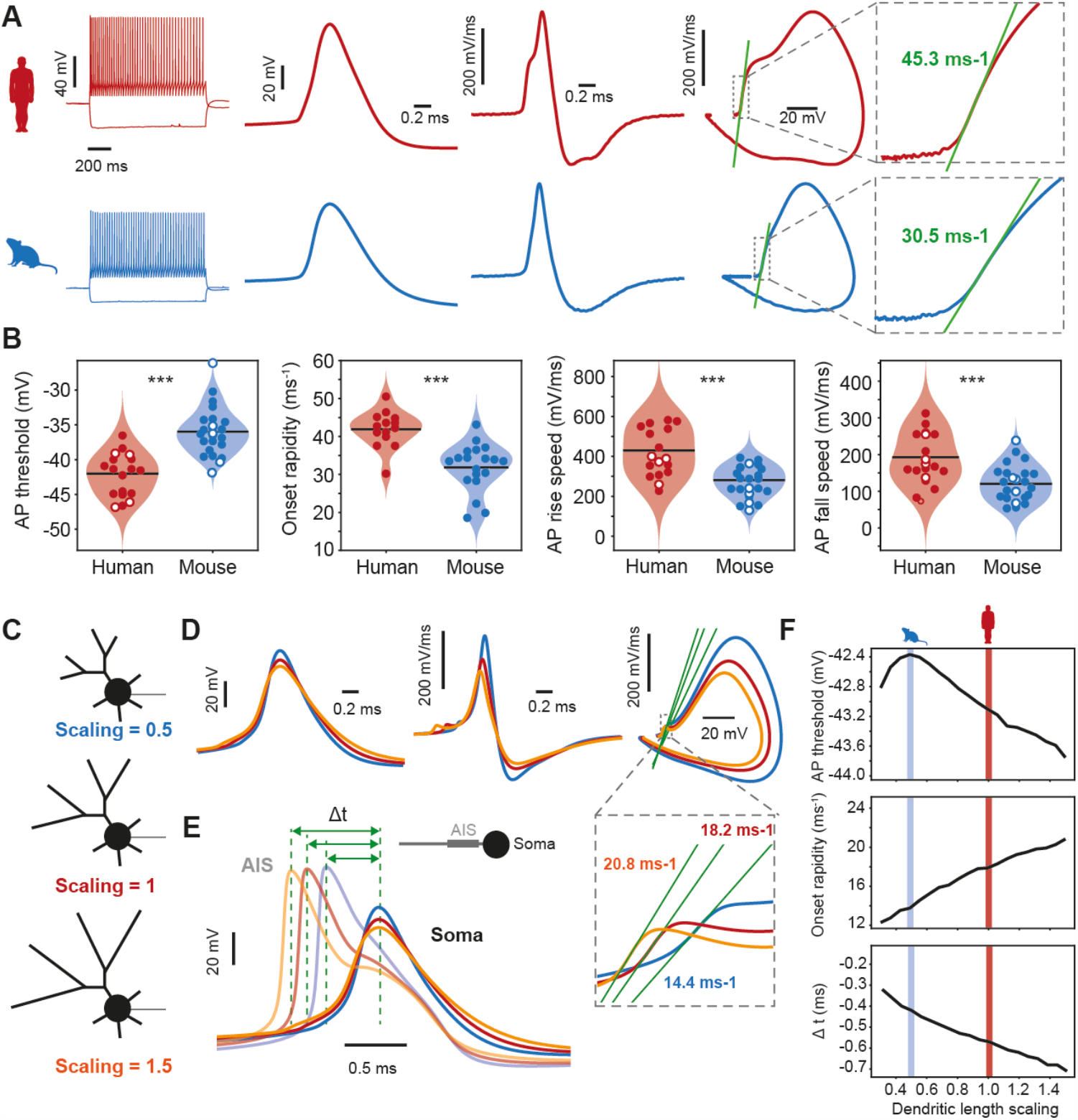
Mechanisms of fast onset kinetics and early AP initiation in AIS in human FSINs. (A) Example AP traces recorded in human (red) and mouse (blue) FSINs. From left to right: AP firing in response to a long square current injection, AP waveform, AP derivative, AP phase plot, onset rapidity (slope fit of the phase plane). (B) Cross-species analysis of AP parameters. From left to right: AP threshold p= 4.9 × 10^-7^, t-test. Onset rapidity p=2.2 × 10^-5^, t-test. AP rise speed, p=0.0001, t-test. AP fall speed, p=0.0003, t-test. Sample size: human n=18 neurons: 14 our lab (filled circles), 4 from AIBS (open circles); mouse n=24 neurons: 19 our lab (filled circles), 5 from AIBS (open circles). (C) Three human models were generated: one with normal dendritic length (scaling=1, red), elongated dendritic length (scaling=1.5, orange) and shorter dendritic length similar to mouse morphology (scaling=0.5, blue). (D) Up- and downscaling of the dendritic length in human model results in a lower threshold, faster onset rapidity and earlier AP initiation at AIS relative to the soma. Example traces from left to right: AP waveform, AP derivative and AP phase plot, and onset rapidity (gray square). (E) Example AP traces in AIS and soma are shown for three models: longer dendritic length leads to earlier AP initiation in AIS relative to soma. (F) Upscaling of dendritic length leads to more negative AP threshold, faster AP initiation kinetics (onset rapidity) and earlier AP initiation in AIS relative to soma, shaded areas indicate dendritic length that corresponds to human (scaling =1, red) and mouse (scaling = 0.5, blue) neurons.

Theoretical and experimental studies point to dendritic size, or dendritic impedance load, as one of the critical parameters influencing fast AP onsets (*27*–*29*). Therefore, we tested how dendritic size would affect AP threshold and onset rapidity in FSIN models. We used a model based on human morphology as a starting point and created two additional models with the size of dendrites scaled to 0.5 of the initial length and a model with the dendritic size scaled to size to 1.5 of the initial length. We then simulated APs in these 3 models both in soma and in the axon initial segment (AIS). We find that scaling up dendritic length decreases AP threshold and increases onset rapidity, and results in an earlier AP generation in the axon relative to the somatic AP, while scaling down the dendritic size leads to the opposite effect (Fig. 5C-G). This indicates that longer dendrites of human FSINs contribute to faster AP initiation, that might be even faster in AIS relative to the soma where APs are experimentally measured.

### Fast synaptic output in human FSINs

The ultimate output function of FSINs is fast inhibition of predominantly pyramidal neurons. To better understand whether human FSINs can generate fast synaptic outputs, we performed whole cell recordings in synaptically connected pairs of FSINs and pyramidal neurons in L2/L3, where the FSIN was presynaptic to a nearby pyramidal neuron. The pyramidal neuron was filled with CsCl-based internal, that amplified inhibitory postsynaptic responses and resulted in large unitary negative currents (outward Cl^-^ currents) in pyramidal neurons in response to activation of FSINs. Such amplification of responses allows to precisely determine the onset latency of responses on an event-by-event basis. We evoked trains of APs in the FSINs and determined the amplitude and timing of unitary postsynaptic currents (PSC) in the pyramidal neuron relative to the peak time of the presynaptic AP (Fig. 6A). We recorded 1800 events from 15 human pairs and 2400 events from 20 mouse pairs that showed reliable responses and only in 5% of events (93 in human and 157 in mouse neurons) there were failures in response that we excluded from further analysis.

**Figure 6.**
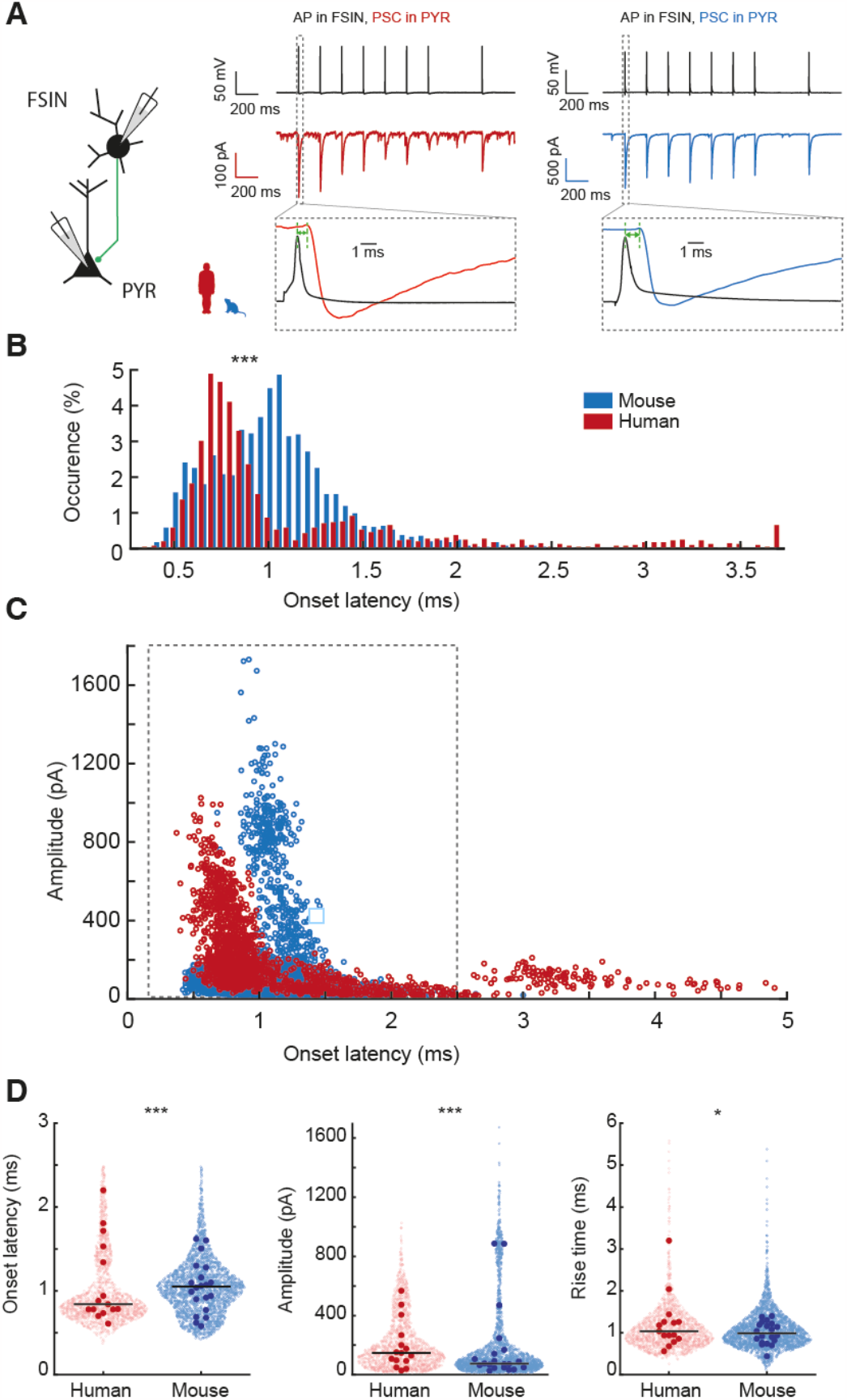
Fast synaptic output of human FSINs to pyramidal neurons. (A) Example traces of paired recordings of FSIN to pyramidal connections performed in mouse and human neurons. APs in FSINs resulted in negative postsynaptic currents (PSCs) in connected FSINs. (B) Distribution of onset latencies of unitary PSC events for human and mouse pairs shows faster PSC onset latencies in human pyramidal neurons, ***P=2.6 × 10^-46^, two-sample Kolmogorov-Smirnov test. (C) Onset latencies of individual responses in FSINs are plotted vs their amplitudes. Events from pairs with monosynaptic connections have onset latencies < 2.5 ms (framed in gray). Sample size: human n=1707 PSCs from 15 recorded pairs; mouse n= 2243 PSCs from 20 pairs). (D) Onset latencies, amplitudes and rise times of only monosynaptic connections (framed in gray in B) are shown as individual events (open circles) and as median values for each connection (filled circles), black horizontal lines are median per recorded connected pair. Statistics individual events: onset latencies ***p=4.3 × 10^—21^; amplitudes ***p=2.3 × 10^-43^; rise times: *p=0.03, MWU test. Median values for each connected pair: not significant, MWU test. Sample size: human n=1569 PSCs from 15 recorded pairs; mouse n= 2224 PSCs from 20 pairs).

We first asked whether human and mouse FSIN inhibitory connections to pyramidal neurons differed in their synaptic delays – onset latencies of the synaptic responses. The distributions of onset latencies were indeed significantly different. However, only in human connected pairs we found high variability in synaptic onset latencies that ranged from 0.5 ms up to 4 ms: most events were very fast (∼0.7 ms), while a small fraction of events had onset latencies longer than 2.5 ms (Fig 6B, D). When plotted against their amplitudes, onset latencies revealed a sharp separation for events with higher amplitudes, while events with extremely long onset latencies were small. Such small events with longer synaptic delays might arise from inhibitory synapses at very distant dendritic locations on pyramidal neurons or might be disynaptic instead of monosynaptic connections. Complex cascades of such multisynaptic events were previously observed in human but not mouse cortex and depend on both inhibitory as well as excitatory action of GABA (*9, 30*). Regardless of their origin, these events were not present in mouse FSIN to pyramidal pairs and we thus focused our analysis only on the events with onset latencies within the monosynaptic range: <2.5 ms. For these events, FSIN to pyramidal synapses were faster and generated response in pyramidal neurons with shorter latencies (median [Q1-Q3] = 0.84 ms [0.72-1.15]), while in mouse synapse the median onset latency was longer (1.03 ms [0.82-1.21]). When comparing the most frequently occurring values of onset latencies in all events - the mode of the distribution, human inhibitory synapses were even faster (mode=0.74 ms) than those of mouse (mode=1.06 ms), indicating that most common FSIN synaptic output in human cortex is a fraction of a ms faster despite potentially longer axonal and dendritic paths. Moreover, human PSCs had larger amplitudes compared to those of mouse (human=155 pA [82-306], mouse= 84 pA [50-169], Fig. 6D) with slightly slower kinetics (10 to 90% rise times: 1 ms [0.78-1.3] in human and 0.96 ms [0.76-1.24] in mouse). When analyzed as median values for all events per connection, this variability is partially lost, and the median onset latencies, amplitudes and rise times are not significantly different between mouse and human pairs, although their median values remain in the same range (Onset latency: median [Q1-Q3], human: 0.84 ms [0.78-1.48], mouse 1.05 ms [0.83-1.22]; PSC amplitude: human: 147 pA [100-250], mouse=74 pA [49-155]; Rise times: human: 1.04 ms [0.87-1.26], mouse: 0.99 ms [0.79-1.2]; sample size: human n=15 connected pairs, mouse n=20).

These results show that human FSINs are able to generate faster and stronger inhibition of target pyramidal neurons.

### Kinetics of disynaptic inhibition through FSINs

Cortical computation relies on feed-forward and lateral inhibition between excitatory neurons through inhibitory FSINs. Simultaneous recordings of two pyramidal cells connected through FSINs show disynaptic delays of 3.72 ± 0.27 ms in human cortex (*15*). From the experimental and modeling data we obtained here, we can now deconstruct the overall disynaptic inhibition latency into its mechanistic components that underlie the synaptic input-to-synaptic output transfer function. Figure 7 shows the step-by-step temporal delays from excitatory synaptic input onto FSINs to inhibitory response in target postsynaptic pyramidal neurons in disynaptic loops in the human and mouse cortex. Firstly, we experimentally recorded similar time delays between AP generation and EPSP initiation in synaptically connected pyramidal cell to FSIN pairs from human and mouse cortex (human=1.31 ms, mouse=1,14 ms, Fig. 2C). Secondly, our findings in models show that the two-fold stronger excitatory synapses and fewer dendrites in human FSINs can accelerate EPSP transfer to soma and compensate for the longer human dendrites, resulting in similar time delays of 0.85 ms (Fig. 4H). Next, we observed faster AP initiation kinetics in our experiments (Fig.5B) and earlier AP initiation in AIS relative to soma in human FSIN model (human=0.6 ms, mouse=0.4 ms, Fig. 5H). Finally, our experimental results in FSIN to pyramidal cell connected pairs show that inhibitory output is faster in human FSINs (human=0.75 ms, mouse=1.06 ms, Fig. 6B,C). In summary, the morphological and functional properties identified above result in a very similar overall input-to-output speed between species: the total delay of the disynaptic loop is 3.5 ms for human and 3.45 ms for mouse circuits (Fig. 7). Thus, the structural and physiological mechanisms of human FSINs we identified in this study counteract the longer dendritic and possibly axonal paths and lead to conserved fast cortical inhibition in the human cortex.

**Figure 7.**
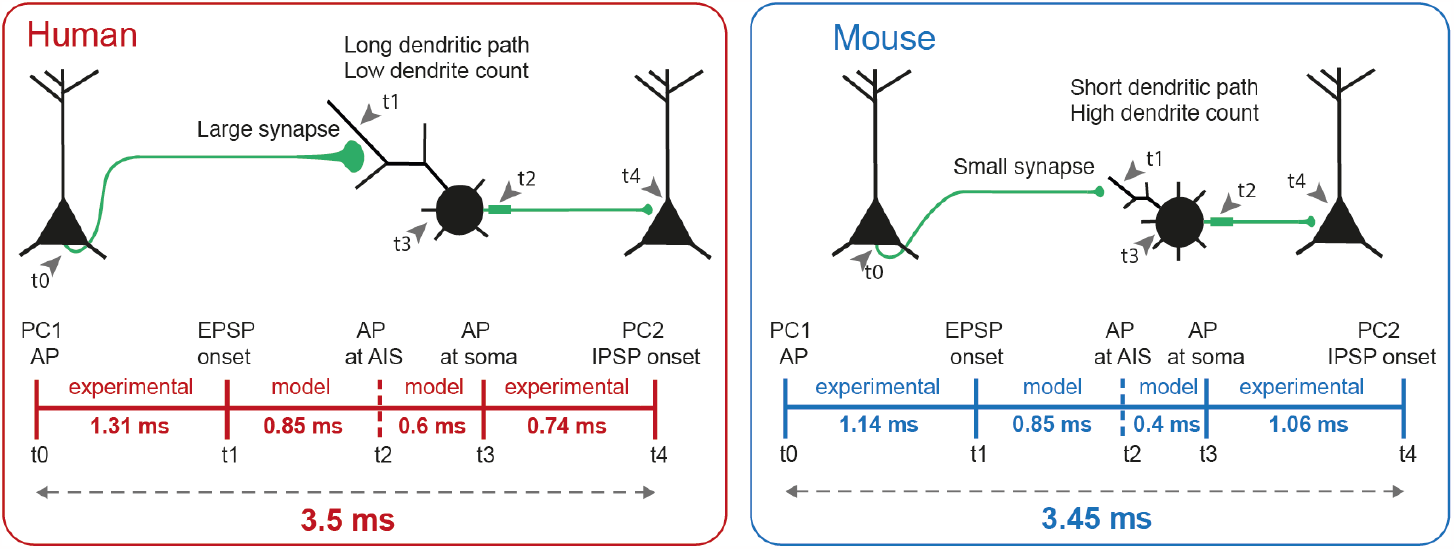
Estimation of the total input-output delay of the fast disynaptic loop through FSINs. Top: schematic illustration of the loop and time series of events from input to output. Bottom: time lines of median delays obtained from experiments and models in this study. Estimated total disynaptic delay is 3.5 ms for human and 3.45 ms for mouse.

## Discussion

In this study, we provide a complete characterization of the input-to-output function in human FSINs. We reveal biophysical mechanisms by which large human FSINs convert distal synaptic inputs to AP outputs as fast as the smaller mouse FSINs, despite two-times longer dendritic paths. Moreover, we show how human FSINs inhibit neighboring pyramidal neurons more rapidly. Combining paired-recordings, digital reconstruction with detailed biophysical modeling we uncovered specific morphological and physiological features that can explain these fast properties: fewer number of dendrites in human FSINs as well as larger synaptic size of human pyramidal to FSIN synapses compensate for the increased dendritic size and boosts the transfer of excitatory synaptic inputs to FSIN soma. In addition, an increased dendritic length facilitates faster output: AP initiation and kinetics are faster in human FSINs and longer dendrites in our models lead to faster AP generation in human AIS. Thereby, large human FSINs can generate fast feed-forward and feed-back inhibition of neighboring pyramidal neurons. For both morphometric and physiological experiments, the experimental data recorded in our lab were combined with publicly available data from Allen Institute for Brain Science (*31, 32*) that support our conclusions and increase the level of confidence and reproducibility of our study.

Our results are consistent with the timing of disynaptic inhibition experimentally recorded in human cortex (*15*). Fast inhibitory loops through FSINs help shape network dynamics and are critical in maintenance of high frequency network oscillations in the gamma range (25-100 Hz). In rodents, gamma oscillations were shown to be suppressed or driven by selective optogenetic manipulation of FSINs in vivo (*33, 34*), supporting the causal role of these interneurons in generation of high frequency network rhythms. In sensory cortex, FSIN-induced gamma activity improve processing of sensory inputs (*34*), while in prefrontal cortex it has pro-cognitive effects and improves goal-directed behavior (*35*). The critical role of FSIN-induced inhibition in cognitive and sensory function might be mediated by the reduction in circuit noise and amplification of signals by these interneurons (*33*). Finally, in humans FSINs are necessary for healthy cortical function, and abnormalities in human FSINs were shown to impair cognitive control and cause cortical dysfunction in schizophrenia (*36*).

Our findings point to several mechanisms that ensure high speed of input-output function in human FSINs. One of these mechanisms is the reduced dendritic complexity of human FSINs compared with mouse FSINs. These results are in contrast to findings in human pyramidal neurons, where dendritic size and complexity are three-fold larger compared with rodents and macaque (*18*). Dendritic complexity in pyramidal neurons is indicative of computational power (*37*) and is increased in high-order associative regions (*38*). A smaller number of primary dendrites that we observe in human FSINs might therefore lead to reduced computational power on the single cell level. However, the loss of dendritic complexity in human FSINs is important for the velocity of signal processing and fast cortical synchronization. This loss of complexity could be compensated by an addition of quantitatively more FSINs. Indeed, spatial transcriptomic studies confirm that the proportion of GABAergic interneurons (where roughly one third are FSINS) to pyramidal neurons is much higher in human cortex (1:2) compared to marmosets (1:3) and mice (1:5). (*6, 39*). Thus, more numerous rather than larger and complex interneurons might regulate the activity in vast dendrites of pyramidal neurons to achieve sufficient inhibition.

Another mechanism behind fast and reliable inputs to human FSINs that we propose is the size of incoming excitatory synapses. This helps boost synaptic inputs and reduce the effect of dendritic filtering. Indeed, experimental measurements in human cortex demonstrate that incoming excitatory synapses on human FSINs have larger active zones with more docked vesicles and functional release sites (*7*). As a result, the increased synaptic energy use could be substantial, since glutamatergic recycling and reversal of ion fluxes through postsynaptic receptors consume most energy in the brain (*40*). Our finding that these large synapses are critical for preserving fast input-output conversion indicates that the increased ATP expenditure that comes with larger inputs is a necessary cost: without it, human FSINs would not be able to function.

APs in neurons are caused by the interplay between voltage-gated Na^+^ and K^+^ channel activation and inactivation. In the present study we do not find differences in Na^+^ current properties of human and mouse FSINs. This is in contrast to our previous report of different Na^+^ current properties in human pyramidal neurons (*41*), where Na^+^ current properties indicate higher functional availability of Na^+^ channels. Possibly, the same subtypes of the pore-forming subunits of voltage-gated Na^+^ channels are expressed in FSINs across species. Work in rodents has shown that parvalbumin positive interneurons express Na_v_1.1 in the soma and proximal AIS, whereas expression in AIS and axon are dominated by Na_v_1.6 (*42, 43*).

Despite the strong similarities between Na^+^ currents across species, we did find differences in AP initiation, with human FSINs showing lower threshold and faster AP onset dynamics. Previous studies have shown that morphological properties such as dendritic length can have profound effect on the rapidity of AP onset in pyramidal neurons (*27, 29*). We find that this principle extends to FSINs, and that AP onset rapidity in our models scales linearly with dendritic length. Furthermore, longer dendrites also resulted in lower AP threshold and earlier AP timing in AIS relative to soma. Therefore, it might be a general feature of human neurons, that fast AP initiation is facilitated by the dendritic size and helps process distal inputs on long dendrites.

Lastly, we investigated the timing difference between a somatic AP in FSINs and inhibitory responses in postsynaptic pyramidal neurons. Although the onset times showed high variability, we find many responses in human neurons with extremely short latencies of 0.74 ms and even shorter. One possibility could be that the synaptic output machinery in human FSINs works even faster than in mice, which already has several specializations to support fast output (*44*). Although we cannot exclude this possibility, the timing difference between soma and AIS, that our models predict, provides a likely explanation. Since the soma spikes relatively late compared to the AIS in human models, the shift in ∼0.2 ms in the response latency distribution might be caused by a shift in ∼0.2 ms in relative AIS timing. However, it cannot be excluded that in the living brain several other mechanisms might converge and play an additional role in supporting the fast processing in human FSIN circuits: active dendritic properties, faster axonal propagation, different receptor subunit expression and kinetics or locations of inhibitory synapses on pyramidal neurons. Furthermore, heterogeneity in myelination patterns of human FSINs (*11, 12*) might help increase velocity of signal transduction and compensate for differences in axonal path lengths at specific synapse locations.

In conclusion, using a combination of experimental and modeling approach, we provide a mechanistic explanation of which human FSIN properties explain conserved fast inhibition and can help human FSINs maintain high temporal precision necessary for cognitive function of the human brain.

## Materials and Methods

### Slice preparation

Human cortical tissue from 28 patients (18 females, 6 males, 4 gender not specified, age mean±SD 44±19 years) was obtained upon neurosurgical resection of non-pathological cortical tissue to access deeper lying pathology (28 patients, Diagnosis: Epilepsy=13, tumor=10, unspecified=5; hemisphere: right=15, left = 7, unspecified=6; Lobe: temporal=16, frontal=10, occipital=1, unspecified=1). All patients consented to the use of material for this study. Tissue from 6 patients was obtained at Beijing Normal University, all others at VU University. All procedures were in agreement with the declaration of Helsinki, Dutch license procedures, ethical standards of VU and Beijing Normal University. Upon surgical removal, healthy cortical tissue was stored in carbogenated NMDG solution at 0 °C and transported to the lab. The pia mater was carefully removed using fine tweezers and tissue was placed in a vibratome (Leica, V1200S) to slice 350 μm thick coronal slices to the cortical surface. Immediately after slicing, each slice was recovered for 12 minutes in 34 °C NMDG solution and subsequently kept at room temperature in carbogenated holding solution.

For animal experiments, all experimental procedures were approved by the Netherlands Central Committee for Animal Experiments and the Animal Ethical Care Committee of the Vrije Universiteit Amsterdam (AVD1120020173124). Mouse brains were obtained with from 44 C57BL/6J (RRID:IMSR_JAX:000664) mice (mean±SD age 30±4.6 days, 25 males, 11 females, 8 unknown gender) after anesthesia with euthasol (i.p. 120 mg/kg in 0.9% NaCl) and transcardial perfusion with 10 mL of 0 °C NMDG solution. After brain removal, it was placed in the vibratome and sliced as described above. All procedures on mice were approved by the animal ethical care committee of the VU university

### Solutions

Slicing NMDG solution (in mM): 93 NMDG, 2.5 KCl, 1.2 NaH_2_PO_4_, 30 NaHCO_3_, 20 HEPES, 25 D-Glucose, 5 Na-L-ascorbate, 3 Na-pyruvate, 10 MgSO_4_, 0.5 CaCl_2_. pH was adjusted to 7.3 before addition of MgSO_4_ and CaCl_2_ to 7.3 with ∼10-15 mL of 5M HCL.

Holding solution (in mM): 92 NaCl, 2.5 KCl, 1.2 NaH_2_PO_4_, 30 NaHCO_3_, 20 HEPES, 25 D-Glucose, 5 Na-L-ascorbate, 3 Na-pyruvate, 2 Thiourea, 2 MgSO_4_, 2 CaCl_2_. pH was adjusted to 7.3 before addition of CaCl_2_ to 7.3 with 1 M NaOH.

Recording solution (in mM): 126 NaCl, 2.5 KCl, 1.25 NaH_2_PO_4_, 26 NaHCO_3_, 12.5 D-Glucose, 1 MgSO_4_, 2 CaCl_2_. All 3 external solutions were adjusted to 310 mOsm.

K-gluconate internal solution (in mM): 115 K-Gluconate, 10 HEPES, 4 KCl, 4 MgATP, 0.3 NaGTP, 10 K_2_-Phosphocreatine, 0.2 EGTA, and 5 mg*ml^-1^ biocytin.

CsCl internal solution (in mM) used for paired FSIN to pyramidal neuron recordings: 120 CsCl, 10 HEPES, 10 TEA-Cl, 4 MgATP, 0.3 NaGTP, 10 Na_2_-Phosphocreatine, 1 EGTA, and 5 mg*ml^-1^ biocytin.

### Whole cell recordings

FSINs were recorded between 200-1200 μm (human) or 100-400 μm (mouse) in whole cell configuration using 3-5 MΩ pipettes filled with K-Gluconate internal solution. The resting membrane potential was measured after establishing whole cell configuration in current clamp mode without current injection. The effective membrane potential at rest was similar between human and mouse FSINs: human: -63.06 ± 4.67 mV (Mean ± SD) n=13; mouse: -65.64 ± 4.44 mV n=16; t-test p=0.13. Spiking profiles were obtained by a series of 1s long DC current injections. In case of paired current clamp recordings for EPSP inputs, 1 or multiple potential presynaptic pyramidal neurons were recorded simultaneously or sequentially and probed for synaptic connectivity. Presynaptic APs were evoked with 8 consecutive 2 ms pulses of 2500 pA at 5 Hz and postsynaptic potentials were recorded using a Multiclamp 700B amplifier (Molecular Devices). Postsynaptic currents were low-pass filtered at 4 kHz and digitized at 50 kHz. For spike profile recordings used in AP analysis, no filter was used and the signal was digitized at 500 kHz. For paired voltage clamp recordings shown in Figure 6, CsCl internal was used in the postsynaptic pyramidal neurons. In a subset of these recordings, presynaptic inhibitory neurons were first probed with juxtasomal stimulation in loose-patch configuration before breaking into whole cell configuration. Only in 5% of recorded responses we observed failures in postsynaptic responses that were excluded from the analysis. All whole cell recordings were performed at 34°C. In addition, we have imported and analyzed current clamp recordings of 5 human and 8 mouse connected pyramidal neurons to FSIN pairs from Allen Institute for Brain Science (AIBS) database (*31*) published in (*9, 45*). For the analysis shown in Figure 2 we extracted EPSP parameters from average traces of 5 recorded traces in response to 10 Hz AP train.

### Nucleated patch recordings

Whole cell recordings were established from FSINs identified by their spiking profiles as above. After switching to voltage clamp mode, the cells were voltage clamped at -70 mV potential, while the holding current was monitored and was similar between human and mouse FSINs: human: -199 ± 178 pA (Mean ± SD) n=11; mouse: -145 ± 159 pA n=10; t test p=0.47. The nucleated patches were extracted while applying a light negative pressure in the pipette (-50 to -70 mbar). After 10-15 seconds, the pipette was slowly pulled away from the cell and the access resistance was monitored. Successful nucleus extraction was estimated when resistance was >750 MOhm. Na^+^ currents were isolated pharmacologically, by adding 3 mM AP-4, 20 mM TEA, and 100µM of CdCl2 into the running solution. After wash-in, the nucleus was positioned at ∼100 μm above the slice surface. We used pipettes of 2.5-3.5 MΩ resistance wrapped in parafilm to reduce the pipette capacitance. Capacitive currents were compensated with pipette compensation (5-8 pF) and whole cell compensation 1.8-3.5 pF, Rs ∼3.5 MOhm, residual artifact was calculated based on subthreshold pulses and subtracted. To minimize this risk that the fast activation time constant could be affected by compensation errors we used 70% prediction/correction during series resistance compensation. To detect the rising phase of the fast kinetics of Na^+^ channels and prevent voltage escapes during the experiments, the temperature during the recording was lowered to 25°C. We monitored the effectiveness of capacitance compensation using a test voltage step pulse. The recordings were sampled at 100 kHz. Correction for the liquid junction potential was applied during the analysis of the traces.

### Morphology

During electrophysiological recordings, cells were loaded with biocytin through the recording pipette. After the recordings the slices were fixed in 4% paraformaldehyde and the recorded cells were revealed with the chromogen 3,3-diaminobenzidine (DAB) tetrahydrochloride using the avidin–biotin–peroxidase method. Slices (350mm thick) were mounted on slides and embedded in mowiol (Clariant GmbH, Frankfurt am Main, Germany). Neurons were examined for completeness of their dendritic trees and only neurons without apparent slicing artifacts and uniform biocytin signal were included (n=5 human neurons and n=10 mouse neurons) and imaged under an oil objective at 100x magnification. Stitched z-stack images were scanned and saved using Surveyor software (Chromaphor, Oberhausen, Germany) at a z-resolution of 0.2 μm. The somatic and dendritic morphology of the neurons was traced using Neuromantic (*46*) and stored in SWC file format. Additionally, 11 human and 33 mouse morphologies from FSINs were obtained from the Allen cell types database (*9, 32*). FSINs were selected using the same electrophysiological criteria and SWC files were downloaded from all cells with full dendritic reconstructions.

Morphological features were extracted using custom made Python scripts using the NeuroM package (*47*). For analysis of path length and terminal segment length, terminal ends where the dendrite was terminated by the slicing where distinguished from true terminals and excluded from analysis. To exclude any effect caused by methodical differences between labs on dendritic diameter, we only included Allen data for dendritic diameter comparison.

### Analysis of Na^+^ currents

Artefacts were subtracted from raw Na^+^ currents using a P/N subtraction protocol. The artefacts were obtained from an averaged response to 5 preceding pre-pulses of -40 mV. Then the artefact was scaled to the voltage of the pulse and subtracted from the raw Na^+^ current. The current was filtered at 20 kHz and the amplitude was calculated relative to the steady-state current at the end of the pulse. Time constants of activation and inactivation were obtained using exponential fits as in Wilbers et al 2023. Boltzmann curves were fitted to the data to obtain half-inactivation and half-activation voltages for each cell. The data was only included for analysis when the R^2^ value of the fit was higher than 0.85.

### Spike profile analysis and FSIN criteria

Spike profiles data was analyzed using MATLAB (RRID:SCR_001622). APs were detected as peaks with at least -10 mV height, at least 20 mV prominence and at least 2 ms separation with other peaks. The threshold was calculated as the voltage at which the slope is 5% of the maximal rise slope. For onset rapidity analysis the phase slope was calculated using a linear fit from 6 us before to 12 us after the point where the phaseplane reached a threshold of 40 mV/ms. The rise and fall speed were calculated as the slopes of linear fits between 30% and 70% from threshold to peak. FSINs were selected based on the following criteria: Membrane time constant < 20 ms, Slope of F/I curve >0.2 Hz/pA, input resistance <200 MΩ, ratio between upstroke and downstroke <1.7. These criteria filtered only typical fast spiking profiles and resulted in a selection of 92% of PV+, 19% of SST+ and 18% of VIP+ cells from mouse lines in the Allen cell types database.

### Statistical analysis

All statistical analyses were performed in Python using scipy and statsmodels packages. In case of unidimensional unpaired analysis, unpaired t-test or Wilcoxon rank sum test was performed dependent on outcome of d’Agostino-Pearson’s normality test. In some plots the bootstrapped confidence intervals of the means were visualized using seaborn.

### Computational model

Artificial morphologies were generated with custom Python scripts based on 5 parameters: number of primary dendrites, non-terminal segment length, terminal segment length, dendritic diameter at base, dendritic diameter at terminal. All dendrites were identical and terminal segments were in the 3^rd^ branching order, which was a typical branching order of terminal segments we observed experimentally in both species. Dendritic diameters were linearly decreased from base to terminal. The initial fit of the electrophysiological parameters of the model (see below) was done with the morphological parameters set to averaged experimental values across species, as to prevent any bias towards one of the species. Computational modeling was performed using the NEURON and BluePyOpt frameworks in Python (*48, 49*). The model contained 4 types of conductances: Na^+^, K^+^, HCN and passive leak, for each of which the maximal conductance was set as free parameter for the dendritic, somatic and axonal compartments. One additional free parameter was used to left-shift the axonal sodium channels in order to mimic low-threshold Na_v_1.6 channels. The 13 free parameters were optimized using an evolutionary approach in BluePyOpt for 500 generations and with offspring size of 100 (evolutionary parameters: eta=10, cxpb=0.7, mutpb=0.7). The fitness of each offspring was evaluated by the SD-scaled absolute difference from the pre-defined feature objectives (Table 1).

The parameter set with the highest fitness (lowest deviation from feature objectives) was used in all modeling figures (Table 2).

Synaptic inputs were simulated with alpha function conductance with time constant set at 0.4 ms and synaptic reversal potential of 0 mV. Specific membrane capacitance was 0.9 uF/cm2, axial resistance 100 Ohm*cm, reversal potential of passive conductance -75 mV, reversal potential for Na^+^ 64 mV, reversal potential for K^+^ -85 mV, axon diameter 0.2 um, reversal potential for HCN -45 mV.

## Author Contributions

Conceptualization: RW, NAG, HDM

Methodology: RW, NAG, AAG, SLWD, TSH

Software: RW

Investigation: RW, AAG, SLWD, TSH, EJM, JH, VDM, DBH

Formal analysis: RW, NAG, AAG, SLWD,

Funding acquisition: HDM, NAG

Resources: SD, PCdWH, SI, DPN, P v.S, IK, GL, TL, YS, CPJdeK, HDM, NAG

Supervision: NAG, YS, HDM and CPJdK

Visualization: RW, AAG, SLWD, NAG

Writing – original draft: RW

Writing – review & editing: RW, AAG, SLWD, NAG, HDM

## Data and Materials Availability

All data needed to evaluate the conclusions in the paper are present in the paper and/or the Supplementary Materials. Furthermore, the morphology reconstructions data used for the analysis in Figure 1 were partially obtained from publicly available database at https://celltypes.brain-map.org/ as dataset (*32*). Electrophysiological recordings from connected pairs used for the analyses in Figure 2 and spike profiles of FSINs in Figure 5. are publicly available at https://celltypes.brain-map.org/ as dataset (*31*) published in (*9, 45*). The electrophysiology and morphology data collected in this study as well as the computational models are provided at the repository DataverseNL: https://doi.org/10.34894/TONOHA.

