## Supplemental material for "Structural and functional specializations of human fast spiking neurons support fast cortical signaling"

### Supplementary Materials

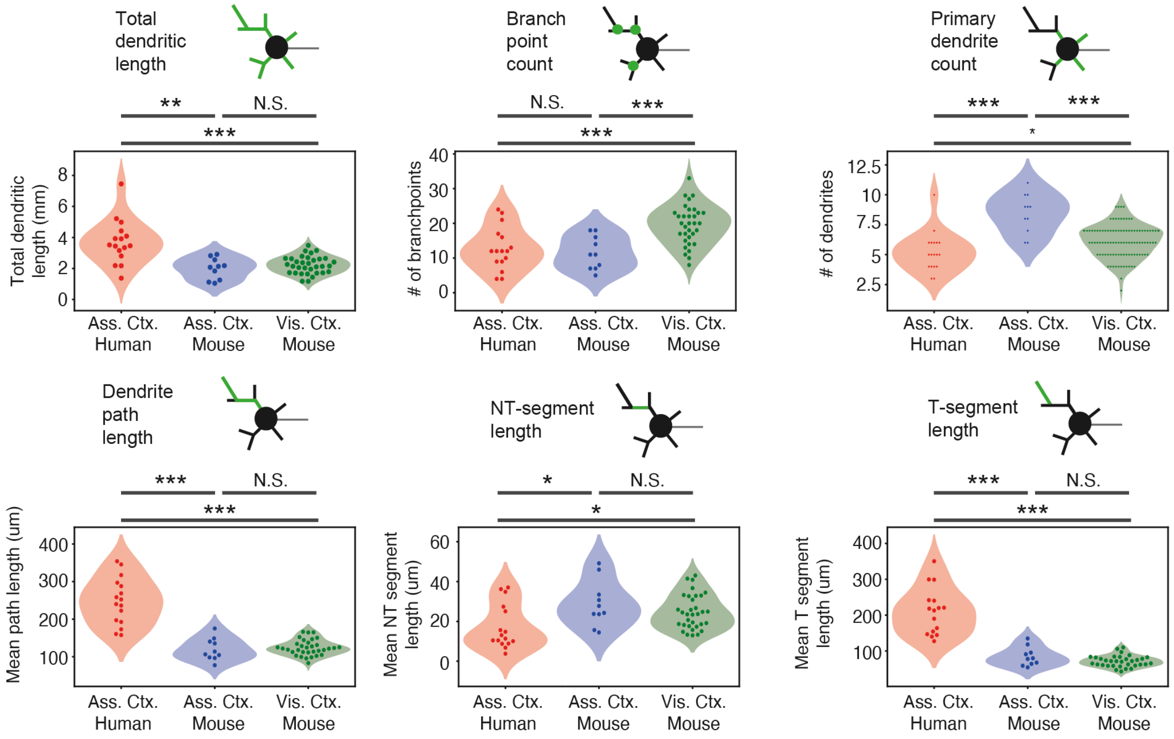


**Supplementary Figure 1. Human and mouse FSIN morphology parameters.** Data are shown separately for the cortical area of FSIN origin in mouse: primary visual cortex and temporal association area. ***$p<0.001$, **$p<0.01$, *$p<0.05$, Wilcoxon rank sum (WRS) test. Human: n=16 FSINs; mouse n=10 FSINs from temporal cortex, n=33 from primary visual cortex.
